# Modeling Reliable Detection Range of Cetaceans Imaged with Infrared Cameras

**DOI:** 10.64898/2026.02.26.708134

**Authors:** Jonathan Bumstead, Casey Corrado Kirsch, Tom Weber, Matt Adams, Carolyn Sheline, Hannah De los Santos, The MITRE Corporation

**Affiliations:** The MITRE Corporation, McLean, Virginia, United States of America; The MITRE Corporation, Bedford, Massachusetts, United States of America

## Abstract

Infrared (IR) imaging systems are used on vessels, platforms, and drones to detect cetaceans several kilometers away, helping to mitigate harm from maritime activities like vessel strikes and pile driving. To ensure operational effectiveness, the reliable detection range (RDR) —the distance at which 100% detection probability is achieved—is a critical metric. This study presents a radiometric model for calculating RDR across a wide range of environmental conditions and system parameters, which enables the evaluation of IR system performance without extensive at-sea data collection.

## Introduction

Cetacean detection using thermal infrared (IR) cameras has been demonstrated with systems mounted on vessels (1), unmanned aerial vehicles (UAVs) (2,3), and stationary structures on shore (4,5). The contrast in images captured with IR systems is based on the temperature difference between the cetacean (e.g., whale blow, dorsal fin, etc.) and background (i.e., the ocean). In addition to image acquisition, IR systems often include some form of object recognition software that labels objects from the video stream (e.g., cetacean blow, ship, etc.), typically requiring a human observer to validate detections (6,7). Several maritime applications can benefit from real-time monitoring of cetaceans that IR imaging provides. For example, these systems can be used to monitor offshore construction activities, where mitigation actions (e.g., stop pile driving) can be taken once a detection is made. Similarly, vessel-mounted systems allow mariners to take evasive maneuvers (i.e., turn, slow down) to prevent vessel strikes, improving safety for both mariners and cetaceans (8,9).

IR systems must deliver cetacean detection results in enough time to successfully perform a mitigation action (e.g., redirect vessel, stop pile driving). Therefore, a critical parameter of these systems is the region over which cetaceans can reliably be detected. Spatial coverage of IR systems is specified by the azimuthal field-of-view (FOV) and the reliable detection range (RDR). RDR is the maximum distance from the camera for which the IR system can detect 100% of cetaceans of interest (10). RDR depends on many factors, but some of the most important include the atmospheric conditions, distance between the target and IR system, the size of the target, and the height of the camera (11,12,4). In foggy conditions for instance, the IR radiation from a whale blow may be too attenuated to be imaged with sufficient signal-to-noise ratio (SNR).

Currently, the RDR of IR systems is determined empirically by generating a probability density function (pdf) of cetacean detections as a function of distance (10,9,4). Assuming a uniform distribution of cetaceans over the sea, several groups have defined the RDR as the distance at which the number of cetaceans detected reaches a maximum (i.e., at the peak of the pdf). Although this empirical method is the current gold standard and ideal for evaluating performance, a large amount of data is required to generate a pdf that accurately represents the system. Further, the RDR is highly dependent on environmental conditions and IR camera parameters, meaning extensive data collection is needed across expected operating conditions to fully characterize the IR system (13). As the use of IR systems for cetacean detection continues to expand, there is motivation to quickly evaluate the performance of IR systems over a range of environmental conditions, cetacean species, camera specifications, camera type (e.g., stationary, rotating, etc.), sensor type (e.g., cooled or uncooled), and more.

Therefore, we developed a theoretical model for calculating useful parameters of IR systems used for detecting cetaceans, specifically the RDR and maximum detection range (MDR). This model has the potential to evaluate a wide range of IR systems in various applications and environmental conditions, creating an avenue to quickly determine the effectiveness of an IR system at detecting cetaceans with sufficient time to perform potential mitigation actions. Although the model could be applied to any cetacean behavior, the results of this paper focus on the detection of whale blows or exhaled breath because the blow’s spatial extent above the sea is a key feature for detection at large distances (14,4). The blow rate is not included in the model, so it does not consider cases for which a whale is at the surface but isn’t spouting (11). Our work is inspired by electro-optical target acquisition modeling that uses radiometry to determine the image quality of the target imaged by an IR camera (15,16,17). From this image quality metric, the probability of object detection can be estimated.

The goal of the work in this paper is to modify the existing electro-optical target acquisition models to include important parameters of IR systems used in cetacean detection such as camera height and orientation, video sequences of events, and blow geometry. First, we describe the developments we made to the IR detection model. Then, the model is applied to an existing IR camera system stationed off the central California coast that detected 1,882 gray whales (4). The robust dataset collected by the system is then compared to the RDR predictions made by our model. Finally, the model is used to conduct several sensitivity analyses that provide insight into which parameters of IR systems most impact cetacean detection performance.

## Materials and Methods

IR systems consist of the optical system, staring array (includes the detector and camera readout integrated circuit), acquisition computer, and detection algorithm (Fig 1) (18). The IR radiation from the target and background propagates through the atmosphere and is collected by the optical system consisting of lenses, an aperture stop, and spectral filter. The camera’s staring array converts the IR radiation to an electrical signal that is read-out, amplified, and processed to generate images that provide temperature information. IR image sequences are then saved or streamed to an acquisition computer with a detection (i.e., object recognition) algorithm that can determine if a cetacean is present, as well as its approximate location. Beyond factors tied to animal behavior, the probability of successful detection depends on a number of system-related factors (e.g., image quality, training data, algorithm type).

**Fig 1.**
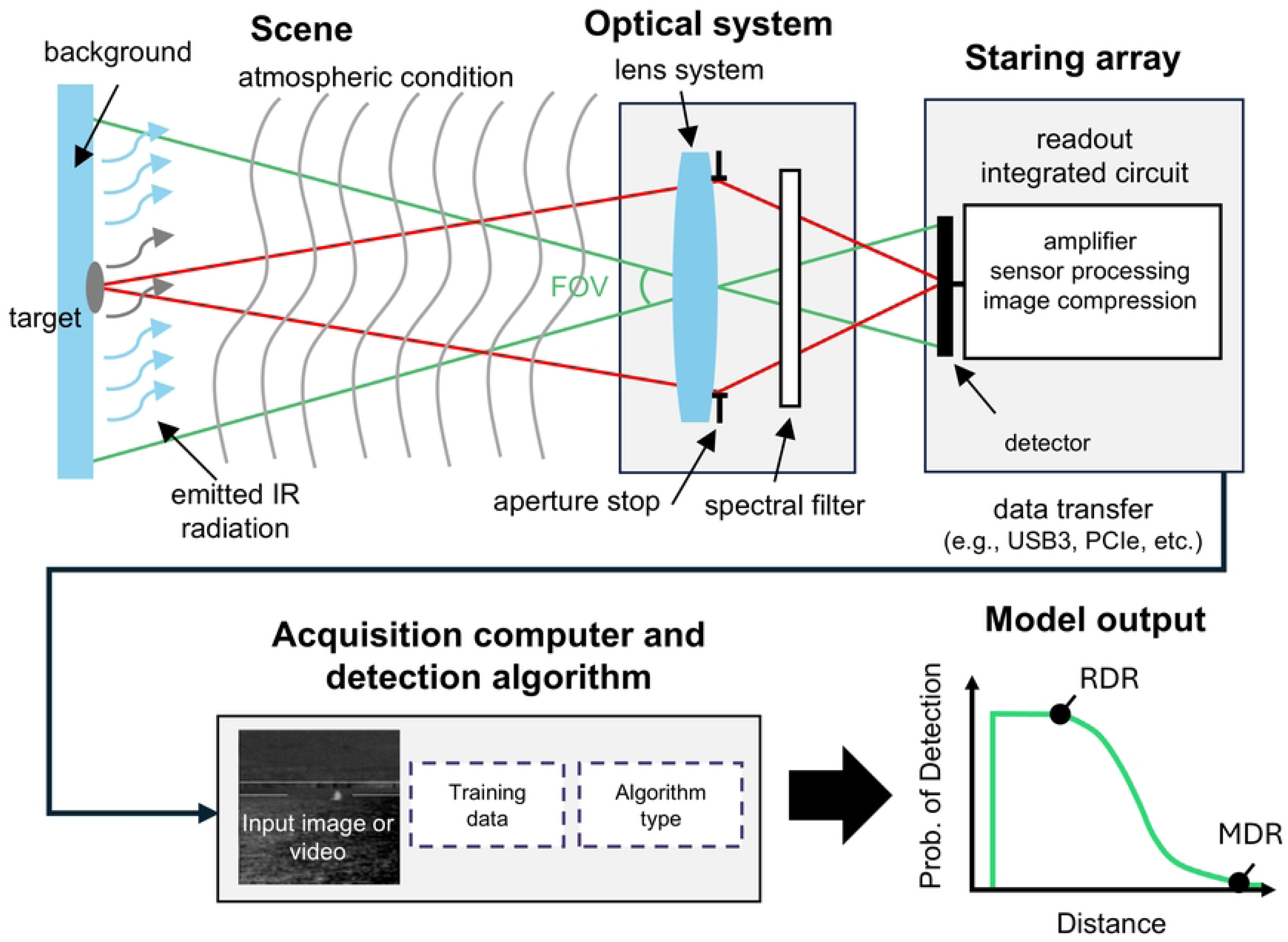
Diagram of IR system from IR emission to detection algorithm output. A single lens represents the multiple lens system typically used in IR cameras. The spectral filter isolates the wavelength range for which the system is designed. Green rays represent the full FOV of the system (chief rays), while the red rays represent the light collecting cone of the system (marginal rays).

The distance between an object and an IR system is projected primarily in the vertical direction of the IR sensor. The maximum distance imaged on the sensor is the horizon line, which depends on the height of the IR system. The maximum distance at which a system can detect a target (i.e., RDR) cannot exceed the horizon line. For any practical IR system, the RDR is much less than the horizon distance and depends on the SNR of the target, the spatial sampling of the target, and the detection algorithm (Fig 2).

**Fig 2.**
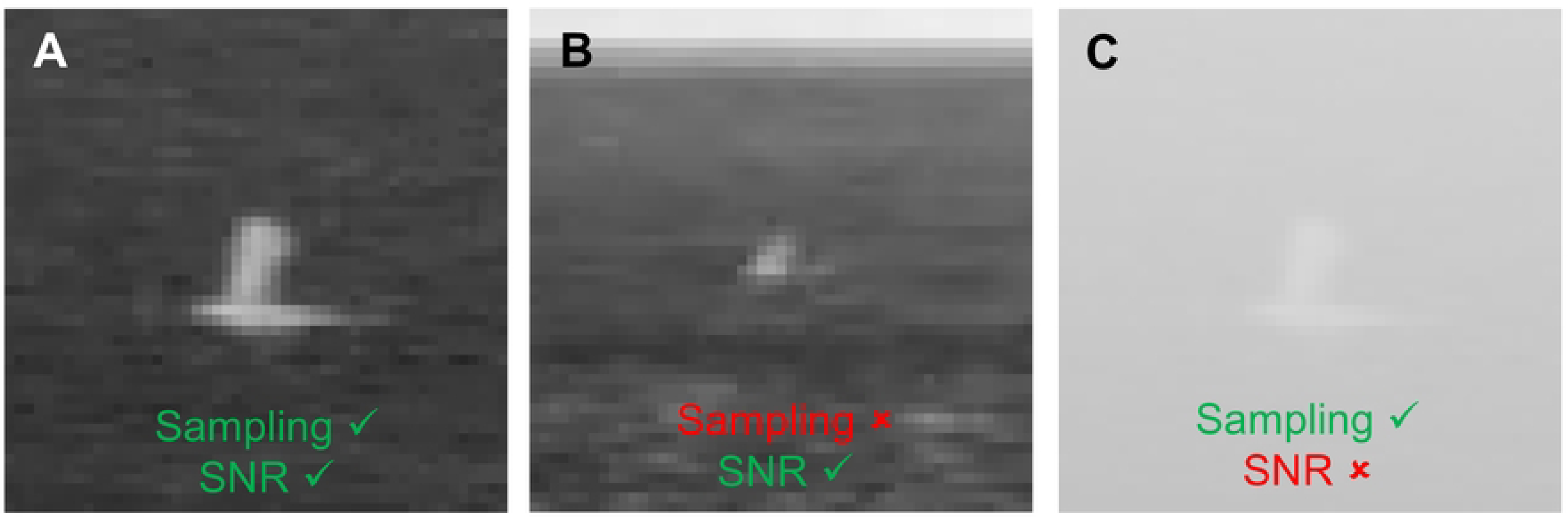
Limitations of reliable detection range due to image quality. (a) Digital zoom of a cetacean blow that may be successfully detected because image has sufficient SNR and spatial sampling of target. (b) Another representative image with worse spatial sampling of a blow that may not be detected by an IR system. (c) Same image as (a) but adjusted to appear as though it was captured on a foggy day. The spatial sampling may be sufficient, but SNR is too low to successfully detect the blow.

The goal of the model presented in this paper is to determine how effectively an IR system can detect a cetacean over its FOV by estimating its RDR. To achieve this, we model the quality of the cetacean image on the sensor using radiometry techniques with a few key modifications in comparison to previous work: the camera elevation relative to sea level, the probability of detection of video sequences, the characteristic dimension of a blow, and model outputs (e.g., RDR) that can be compared to experimental results.

The probability of detecting the cetacean based on the image quality is determined using the targeting task performance (TTP) metric (15). Both the SNR and spatial sampling of the cetacean determine the image quality and therefore the effectiveness of object recognition algorithms. As the FOV increases, objects that are far away may be too pixelated to differentiate on the sensor (Fig 2B). An overview of all inputs and outputs of the model developed here is also provided in the S1 Appendix.

### 2.1 Image quality and targeting task performance metric

Here we briefly describe the signal pathway from IR photon to image contrast and the key takeaways. The signal begins with the thermal emission of IR radiation by the target and background, which can be described using Planck’s law. As the IR radiation propagates, some of it will be absorbed and scattered by the atmosphere between the target and the IR camera. The Beer-Lambert law can be used to estimate the fraction of radiation reaching the camera by converting atmospheric visibility into the attenuation coefficient using Koschmieder’s Law:

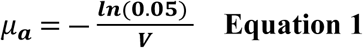

where μ_a_ is the attenuation coefficient and V is the visibility in km (19). Table 1 provides a few example conditions.

**Table 1.** Typical visibility and attenuation coefficients for clear, hazy, and foggy days.

| Description | Visibility (km) | Attenuation<br>coefficient (1/km) | Transmission<br>per km |
| --- | --- | --- | --- |
| Clear | 50 | 0.06 | 0.94 |
| Haze | 3 | 1.0 | 0.37 |
| Fog | 0.5 | 6.0 | 0.0025 |

The IR radiation is collected by the lens system and focused on a detector that converts photons into an electrical signal measured in electrons per pixel. This electrical signal relative to the background and noise terms provides the SNR of the cetacean imaged by the IR system. The overall image quality depends on the SNR calculated using radiometry, as well as the spatial sampling of the cetacean on the sensor (Section 2.5 and S2 Appendix). An object cannot be detected if it is too pixelated on the sensor.

An important component to detecting cetaceans with IR systems is the method for determining the presence of a cetacean in the IR image. Marine mammal observers can manually review the IR video stream, but for practical implementations of IR systems, there have been efforts to develop automatic detection algorithms (1,9).Object recognition algorithms are now commonly generated using deep learning techniques, which require training datasets. Their performance is typically evaluated using metrics (e.g., average precision, classification error) derived from the algorithm’s output on a dataset it was not trained on (20).

The goal of our model is to determine the probability an object (e.g., whale blow) is detected given the image quality predicted for an IR system. Because the performance of detection algorithms developed using deep learning is usually determined empirically using test datasets, it is challenging to model the effectiveness of an object recognition algorithm analytically. Therefore, our model determines the probability of how well a human would be able to identify an object given the calculated image quality, which can be determined based on analytical models (16,15) Deep learning techniques are capable of exceeding human capability in some object recognition tasks, so using the performance of the human visual system (HVS) to evaluate performance may be conservative (20). The HVS response is characterized using a contrast threshold function. Both the imaging system modulation transfer function and the HVS contrast threshold function are used to determine the overall system response for an image collected with an IR system.

The clarity of the object in the image is with reference to the overall system response and can be used to determine how effectively the object can be recognized using several metrics. In this work, we used the TTP metric. Object recognition can be classified into three tasks: detection, recognition, and identification. Detection corresponds to the ability of the algorithm to determine that an object is present relative to a background, without differentiating it from other objects. Recognition corresponds to the algorithm’s ability to differentiate a class of objects (e.g., a vessel vs. a bird vs. a cetacean). Identification corresponds to whether an algorithm can describe the exact type of object (e.g., which type of vessel, type of cetacean). The TTP metric provides the probability that one of these tasks is performed successfully. In this report, all results were calculated for detection of the object.

### 2.2 Characteristic dimension of cetacean blow

An important parameter to calculate the image quality depends on the size of the target and how well it is sampled by the camera sensor. In this work, we focus on detecting whale blows, but the spatial profile of any cetacean could be used in our model. The spatial extent of the blow that is captured by the IR camera is described using the characteristic dimension (CD). CD is equal to the square root of the object area viewed by the camera. Smaller CD results in poorer spatial sampling and therefore worse image quality.

We assumed the volume of the blow could be approximated as a cone or ellipsoid with height h_w_ and radius r_w_, similar to modeling the whale body as an ellipsoid (21). The CD is then calculated by the 2D projection of the blow. In windy conditions, the blow height may be reduced, resulting in a smaller CD. However, our current model does not factor in this effect or any changing CD to the blow over time. The CD for a cone or ellipsoid is given by:

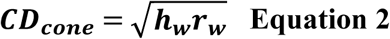

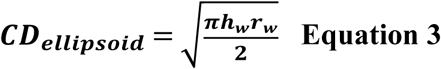

### 2.3 Azimuthal FOV and position

The azimuthal (horizontal) FOV of a stationary IR system depends only on the focal length and the size of the sensor.

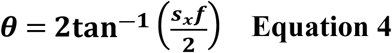

where s_x_ is the sensor size in the x-direction, f is the focal length of the lens, and θ is the full azimuthal FOV (Fig 3A). The azimuthal position of the cetacean θ_target_ can also be calculated from the image using geometric calculation. Assuming that the camera position center is the 0° azimuth position, then the azimuthal position of the cetacean is:

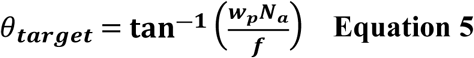

**Fig 3.**
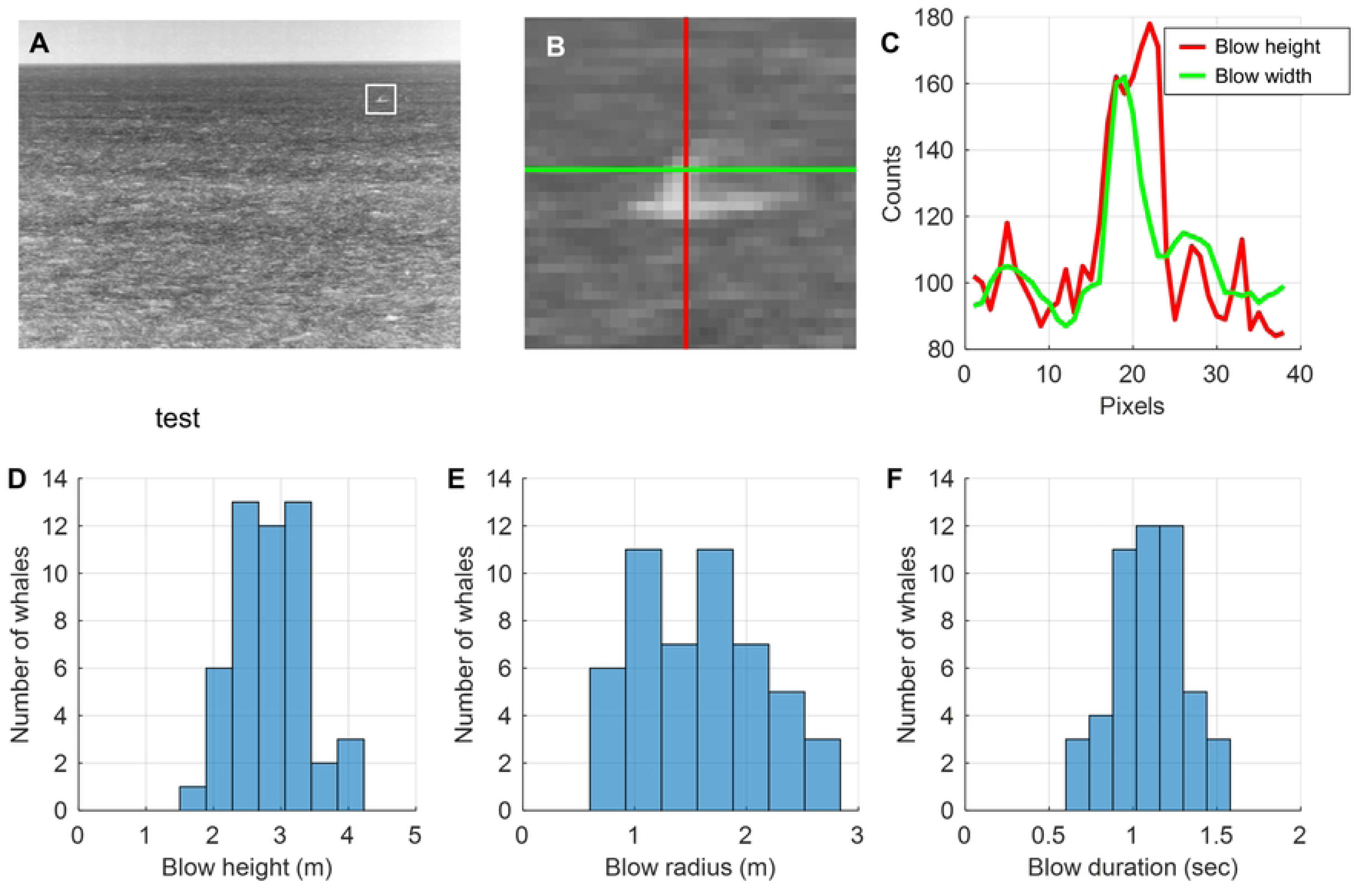
Spatial coverage of an IR camera. (a) The angular FOV set by the focal length and sensor size. (b) Projection of 8×8 pixel sensor onto the sea surface. The gray region is the FOV with each semi-annulus corresponding to a pixel. The blue circles symbolize cetaceans randomly distributed over the sea. (c) The maximum detection range is limited by the distance to the horizon. The red circle marks the position of a whale. The camera height and angle have been exaggerated relative to the curvature of Earth. (d) Representative IR image with whale. The dashed green and red lines correspond to the diagram in c.

where w_p_ is the width of the pixel and N_a_ is the number of pixels in the horizontal direction between the blow and the center of the camera sensor.

Panning IR cameras can be used to extend the FOV in the horizontal direction at the expense of temporal resolution and SNR. Multiple cameras can also be used. A detailed analysis of static and rotating cameras is in Section 3.2. Given the typical IR system configuration, the range is primarily projected onto the vertical direction of the sensor. Therefore, the vertical FOV is better described as a minimum and maximum range (horizon line), which depends on the focal length, sensor size, and camera height.

### 2.4 Range estimation

Once a target has been detected in an image, the position of that target can be calculated using a broad range of localization algorithms that can provide estimates for the object’s coordinates. Deep learning techniques used for sea surface object detection and localization based on image data have become increasingly popular (22). The success of these techniques depends largely on the type of data and the amount of data used as inputs into the algorithm. However, given the typical configuration of IR systems, the cetacean position and range can also be determined using a more straightforward geometric analysis of the image (23).

By calculating how the sea surface is imaged onto the camera sensor, the cetacean range can be determined by measuring the number of vertical pixels between the horizon and marine mammal (Figs 4C-D). It should be noted that the following assumes that the object recognition algorithm successfully detected the cetacean.

**Fig 4.**
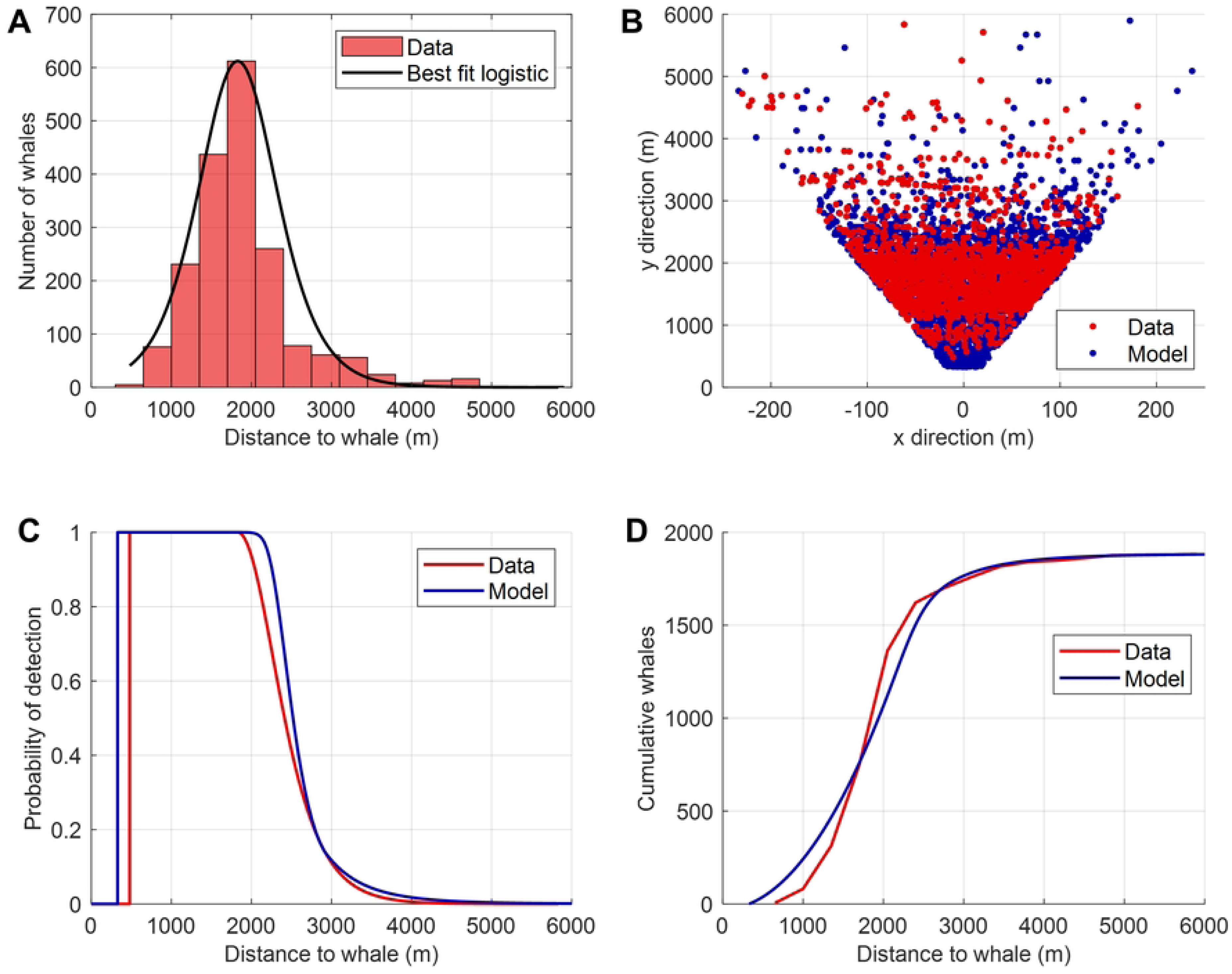
Blow parameters for gray whale dataset. (a) Representative image captured with IR system in Guazzo et al. study. (b) Digital zoom of whale blow with blow cross section marked with red and green lines. (c) IR camera counts for blow cross section in b. (d) Histogram of blow height for 50 whales from study. (e) Histogram for blow radius from study. (f) Histogram of blow duration.

### 2.5 Camera height and spatial sampling

The perspective of the camera due to its height above sea level is a key difference between previous work in object detection performance analysis and cetacean detection with IR systems (15). The horizon distance and RDR increase as the height of the camera increases. Due to the improved perspective, higher cameras have denser sampling over the sea at farther distances (24). However, the pixel size as a function of distance and azimuth is nonlinear. Fig 3C shows a projection of an 8×8 pixel sensor projected onto the sea that accounts for the Earth’s curvature. More sea area is covered per pixel as the distance increases. Because the cetacean is positioned along the surface of the sea, the nonlinear sampling of the sea must be accounted for in the model. The approach used in this work was to scale the effective size of the cetacean according to the pixel projection onto the sea. The pixel projection on the sea was determined using the geometric model (S2 Appendix). The number of pixels per blow was then determined as a function of distance using:

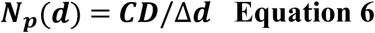

where N_p_ is the number of pixels sampling a blow in one dimension, d is the distance between the camera and the cetacean, and Δd is the radial length of the projected pixel on the sea surface at distance d (S2 Fig).

### 2.6 Probability of detection for video

The model determines the probability of detecting a cetacean in a single image collected by the specified IR system, environmental conditions, and blow characteristics. Given the frame rate and duration of the blow, multiple images are captured by the IR system. The video stream improves the probability of detection. To estimate the probability of successfully completing a task with M images, the following equation was used:

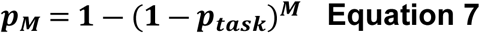

where p_M_ is the probability of completing a task with M images and p_task_ is the probability of completing the task with a single image. Therefore, the duration of the event (e.g., blow) and the frame rate affect the RDR.

### 2.7 PDF and CDF

The model also calculates the expected cetacean detections as a function of distance, the probability density function (pdf), and cumulative distribution function (cdf). The model assumes that the cetaceans are distributed over the sea uniformly with density ρ_c_ (cetacean/km^2^). To determine the number of cetaceans at distance *d*, we need to multiply the cetacean density by the surface area covered by the sensor at distance *d*. Due to the curvature of the earth, the projection of the sensor onto the sea surface is nonlinear. Therefore, the area covered at a distance d is a semi-annulus that increases in area as d increases (Fig 3B). The number of cetaceans N_c_ predicted at a distance d from the camera is then given by:

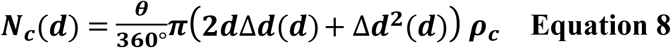

where θ is the full azimuthal FOV in degrees and ρ_c_ is the density of cetaceans at the surface. Plotting this function for a typical IR system provides the predicted number of cetaceans that exist at distance d from the camera. To determine the number of cetaceans detected by the IR system, the number of cetaceans within the FOV needs to be multiplied by the probability of detection function.

## Results

The goal of this work was to develop an IR model that could predict the RDR for IR systems. In this section, we first compare our model to an existing dataset of gray whales. The model can be used for any cetacean with the most important factor being the difference in blow size. The model is then used to conduct a sensitivity analysis to determine the parameters that affect IR system performance (e.g., probability of detection, RDR).

### 3.1 Comparison to existing study

To validate the model, we used data collected in Guazzo’s 2018 study on migrating eastern North Pacific gray whales (e.g., *Eschrichtius robustus*) (4). The onshore IR camera (FLIR F-606, f=100mm, 640×480, 17μm pixel size, uncooled VOx microbolometer) detected 1882 whale blows over four days, which provided a robust sample size for validation. The camera was mounted 28.1 m above sea level. To estimate the performance of the uncooled thermal camera, the dark current density was estimated to be 10^-7^ A/cm^2^, and the read-out noise was set to 500e^-^/pixel/frame.

The blow parameters used in the model were based on video sequences of 50 whales collected with the IR system (Fig 4). Position information and time stamps for each blow were confirmed with a marine mammal observer. The blow height, blow width, and blow duration were then calculated for each blow. Average values for each parameter were then used in the model: blow height = 2.84 ± 0.51 m, blow radius = 1.61 ± 0.54 m, and blow duration = 1.1 ± 0.21 sec.

The distribution of whale blows over the FOV was plotted using the localization information provided from the study and the camera properties (Fig 5A). To provide a comparison to the model, whale blows were assumed to be uniformly distributed over the FOV with a density that matches the detections made in the study. To provide a qualitative visualization of how whales may appear if spotted randomly over the sea, the number of whales predicted at each range was plotted randomly over the FOV (Fig 5B).

**Fig 5.**
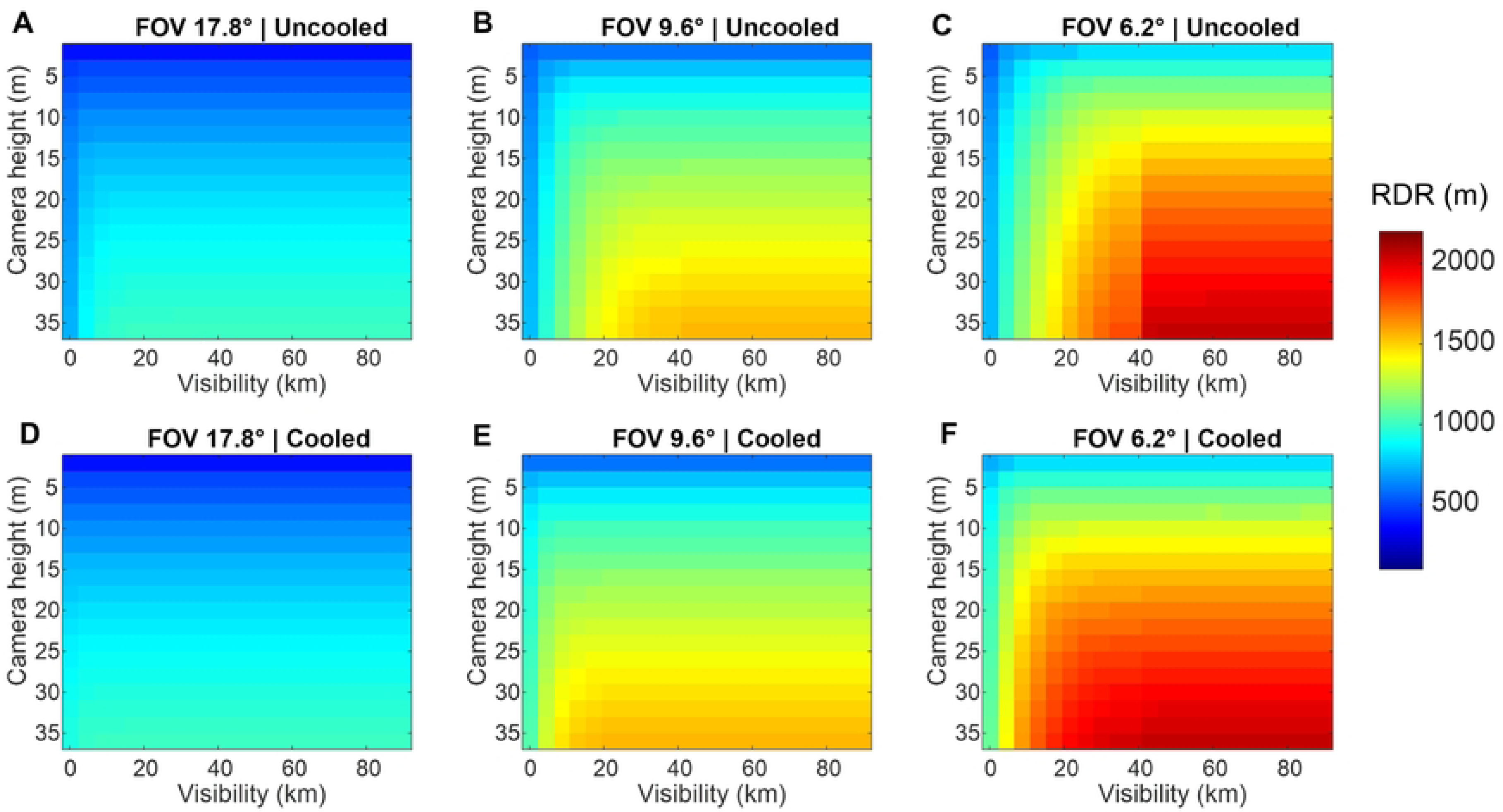
Infrared camera model validation. (a) Histogram of detections as a function of distance from onshore IR system. (b) Distribution of whales over the sea for the experimental measurement (red) and the model results (blue). (c) Probability of detection curve of the model compared to the curve predicted using empirical method. (d) Cumulative distribution for whales detected as a function of distance.

The probability of detection curves and the cumulative IR detections were plotted with the model results (Fig 5C and 5D). The calculated RDR was 1.8km and 2.0km for the empirical method and our model, respectively.

### 3.2 Sensitivity analysis

A major application of the model is to determine how RDR is affected by system parameters and environmental conditions. A sensitivity analysis conducted with the model could inform which IR systems are most capable of detection, as well as the conditions during which a particular system may no longer effectively detect cetaceans. The effort required to evaluate the parameter space experimentally is enormous compared to running a sensitivity analysis with our model.

In this analysis, we attempted to use realistic parameter ranges reported in literature, as outlined in Table 2 (4,12,24). Due to the high dimensionality of the parameter space, the analysis uses several available lenses and sensors instead of sweeping through camera parameters (e.g., focal length, number of pixels, etc.). If not otherwise noted in the results, the default parameter was used in the simulation. For example, plots sweeping camera height and lens type would have all other parameters set to default as shown in Table 2.

**Table 2.** Input parameters used in sensitivity analysis.

|  | Minimum | Maximum | Default |
| --- | --- | --- | --- |
| Camera height (m) | 2 | 36 | 15 |
| Visibility (km) | 0.1 | 90 | 70 |
| Sea temperature (K) | 270 | 309 | 307 |
| Whale blow height (m) | 0.5 | 5 | 3 |
| Whale blow radius (m) | 0.1 | 2 | 1 |
| Whale blow duration (s) | 1 | 5 | 3 |

| Lens label | Focal length (mm) | f-number |
| --- | --- | --- |
| Lens 1 | 25 | 1.1 |
| Lens 2 | 35 | 1.1 |
| Lens 3 | 50 | 1.2 |
| Lens 4 | 65 | 1.25 |
| Lens 5 | 75 | 1.1 |
| Lens 6 | 100 | 1.6 |

**Table 2. Input parameters used in sensitivity analysis.**
| Sensor types | x-dimension (pixels) | y-dimension (pixels) | Pixel size (um) | Frame rate (Hz) | Dark current density (A/cm<sup>2</sup>) | Downstream noise (e-/pixel/frame) | Type |
| --- | --- | --- | --- | --- | --- | --- | --- |
| FLIR F-606 ID | 640 | 480 | 17 μm | 30 | 10<sup>-7</sup> | 500 | Uncooled |
| FLIR FH-313 R | 320 | 256 | 17 μm | 30 | 10<sup>-7</sup> | 500 | Uncooled |
| FLIR A6301 | 640 | 512 | 15 μm | 30 | 10<sup>-12</sup> | 50 | Cooled |

To visualize most of the camera parameters, we generated heat maps of RDR as a function of FOV, camera height, and sensor type (Fig 6). In general, the smaller the FOV and higher the camera height, the longer the RDR, which reflects results in the literature (12,9).

**Fig 6.**
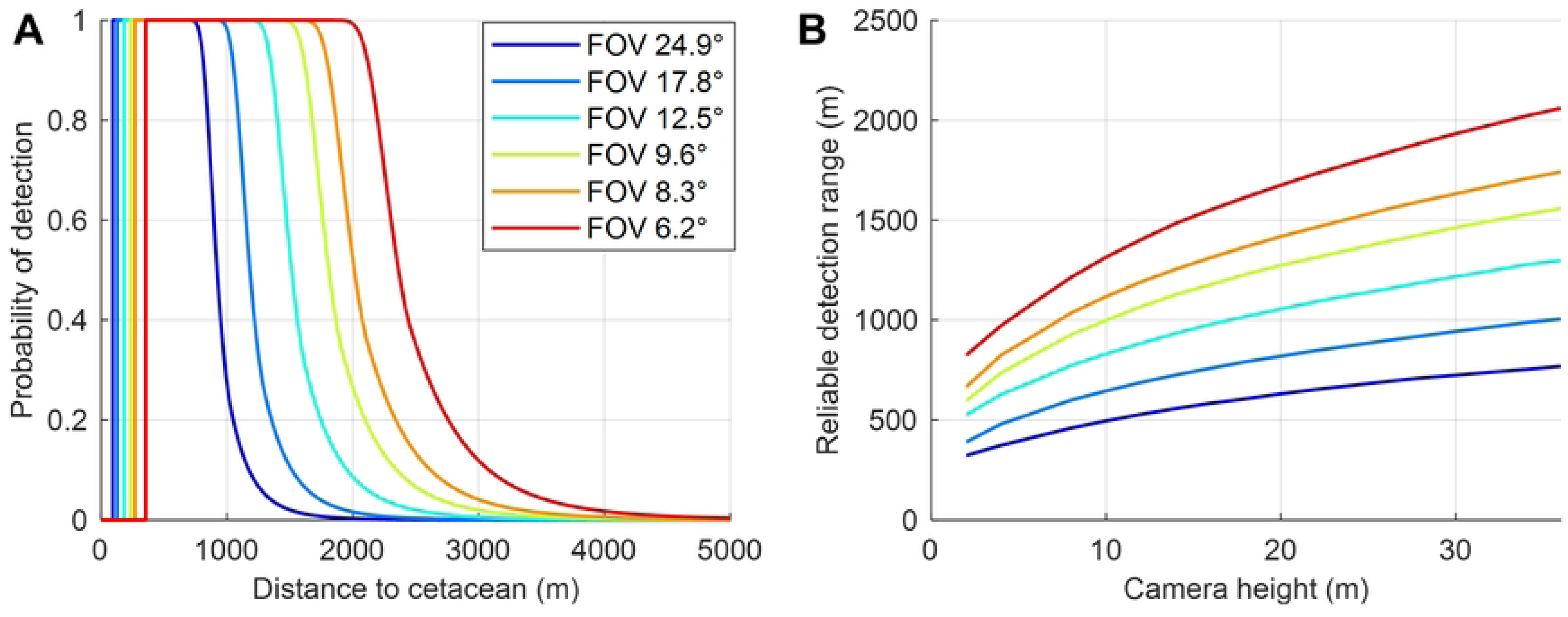
Overview of key parameters that affect RDR. (a-c) RDR as a function of camera height and visibility for an uncooled sensor (FLIR F-606 ID) for three different lenses (Lens 2, 4, and 6 in Table 2). (d-f) Same as a-c, except for a cooled camera with the same sensor size. The temperature of the cetacean and background are 309.15K and 306K respectively.

These plots also reveal the limiting factor for various configurations. For example, Fig 6A and 6D show instances in which the RDR is relatively constant with respect to visibility. The large FOV of the system limits the RDR because of the poor spatial sampling of the image of the cetacean on the sensor. In other words, poor spatial sampling limits the RDR before atmospheric attenuation has any effect on performance. Even for systems with smaller FOV (i.e., higher spatial sampling of cetacean), there is a point at which improved visibility no longer results in increased RDR (e.g., around 40km visibility for the simulation in Fig 6C). Despite the improved visibility, the image quality is limited by the spatial sampling of the cetacean.

Because FOV and camera height play such a critical role in IR system performance, the probability of detection as a function of distance was plotted for various FOVs and camera heights (Fig 7A). For each probability curve, we also calculated the RDR and plotted it for the various conditions (Fig 7B).

**Fig 7.**
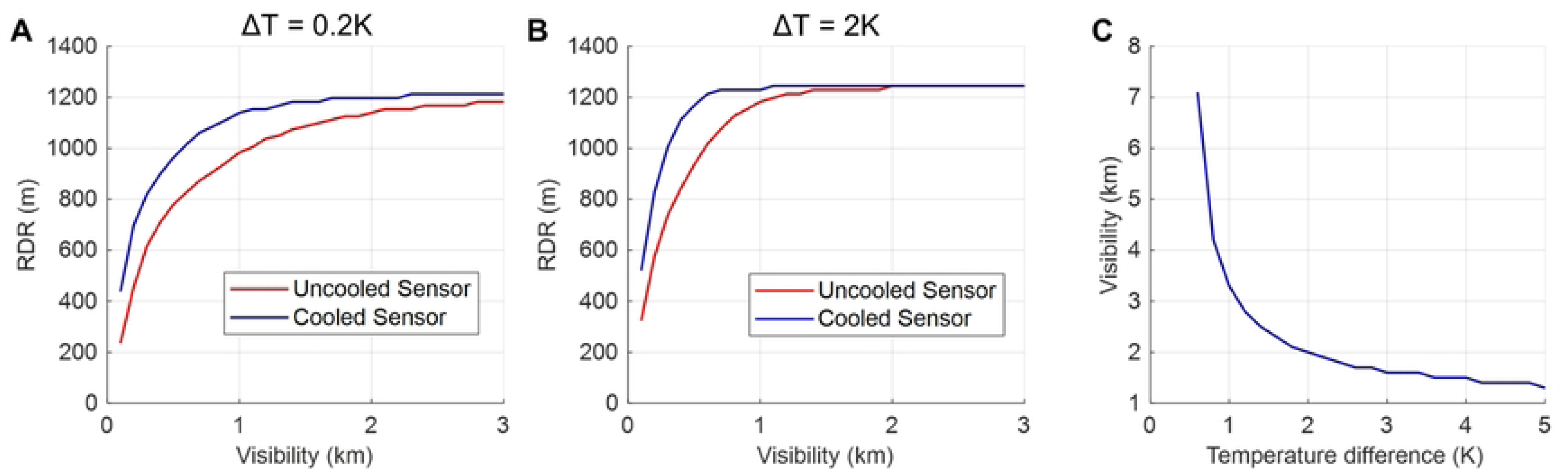
Effect of camera height and FOV on reliable detection range. (a) Probability of detection as a function of distance for a camera height of 15m. The detector parameters are the same with the FOV changing due to different lens focal lengths. (b) Reliable detection range plotted as a function of camera height for different FOV.

To determine the expected improvement with cooled IR sensors, the model was run to calculate RDR for various temperature differences and visibilities (Fig 8). As the visibility decreases, the RDR becomes limited by the SNR of the sensor. In these cases, the reduced dark noise of cooled sensors improves the SNR and, by extension, the RDR. The improved RDR for cooled sensors is greater for cases when there is a smaller temperature difference between the background and cetacean (comparing Fig 8A with 8B).

**Fig 8.**
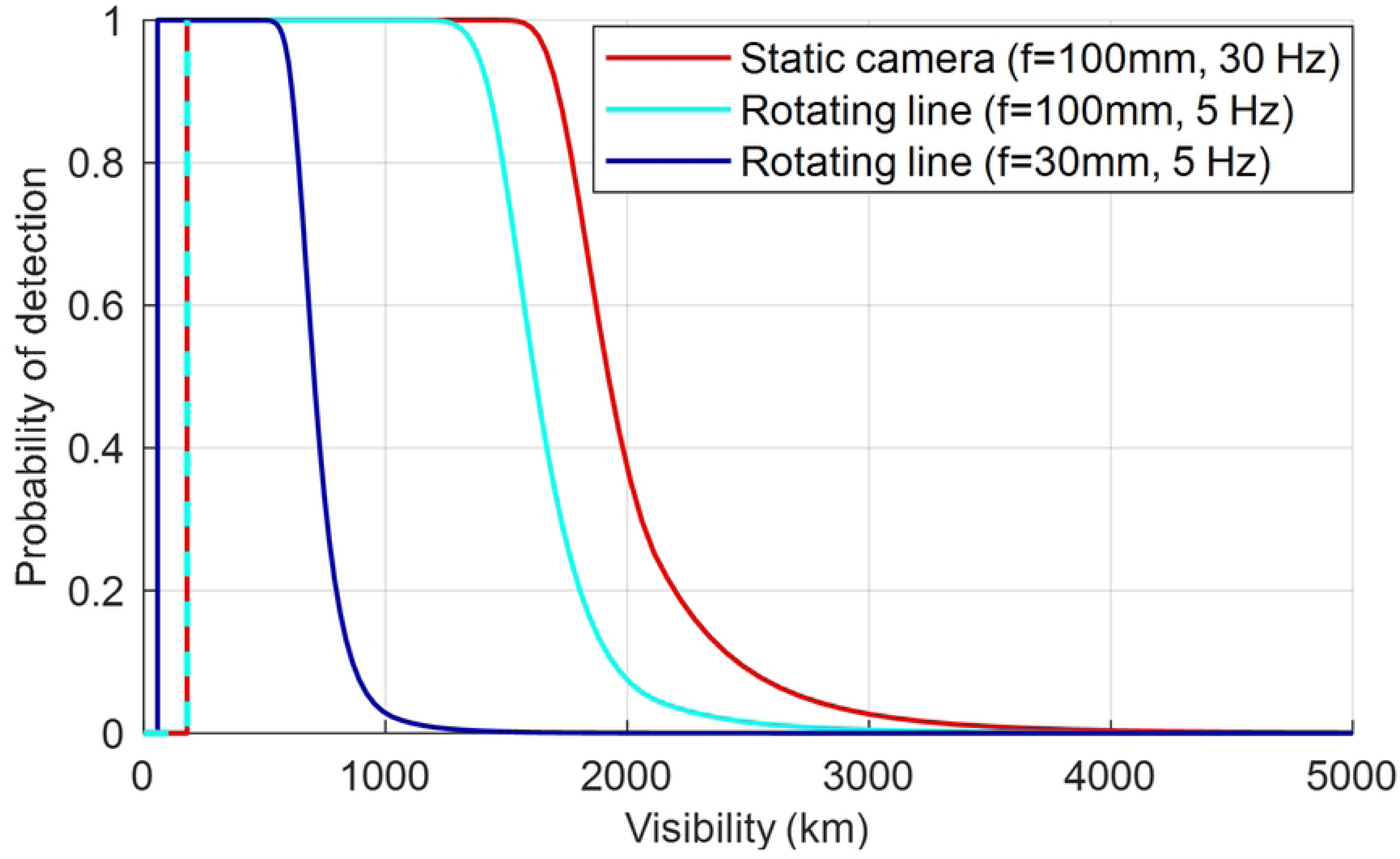
Detection improvement using cooled IR sensors. (a) Reliable detection range as a function of visibility for a cooled and uncooled camera when the cetacean and background temperature difference is 0.2K. (b) Same as b, but for when the temperature difference is 2K. (c) Plot of the visibility at which RDR is the same for cooled and uncooled sensors as a function of temperature difference.

In all cases, the SNR eventually is not the limiting factor, and the RDR is limited by the spatial sampling of the cetacean image on the sensor. Once both the cooled and uncooled sensors reach this point, there is no difference in the RDR achievable by a cooled and uncooled sensor. The visibility at which this occurs was plotted as a function of temperature difference (Fig 8C).

Concurrent ocean coverage (COC) defines the total FOV of all the camera(s) within an IR system. For stationary cameras, multiple cameras can be placed side-by-side to essentially combine their FOVs, resulting in a larger region of COC. It should be noted that camera configuration could result in blind spots and should be checked to ensure there are no coverage gaps. The field-of-regard (FOR) is the entire viewable area of the camera over its rotation range, which is greater than the FOV of the camera when it is stationary. While panning cameras may move slowly, rotating line scanners are capable of 360-degree COC at several rotations per second.

Currently, our model captures the difference in performance between a rotating line scanner and static camera by only changing the frame rate. Due to the fewer images collected per cetacean event, the probability of detection decreases when using a rotating line scanner (Fig 9).

**Fig 9.**
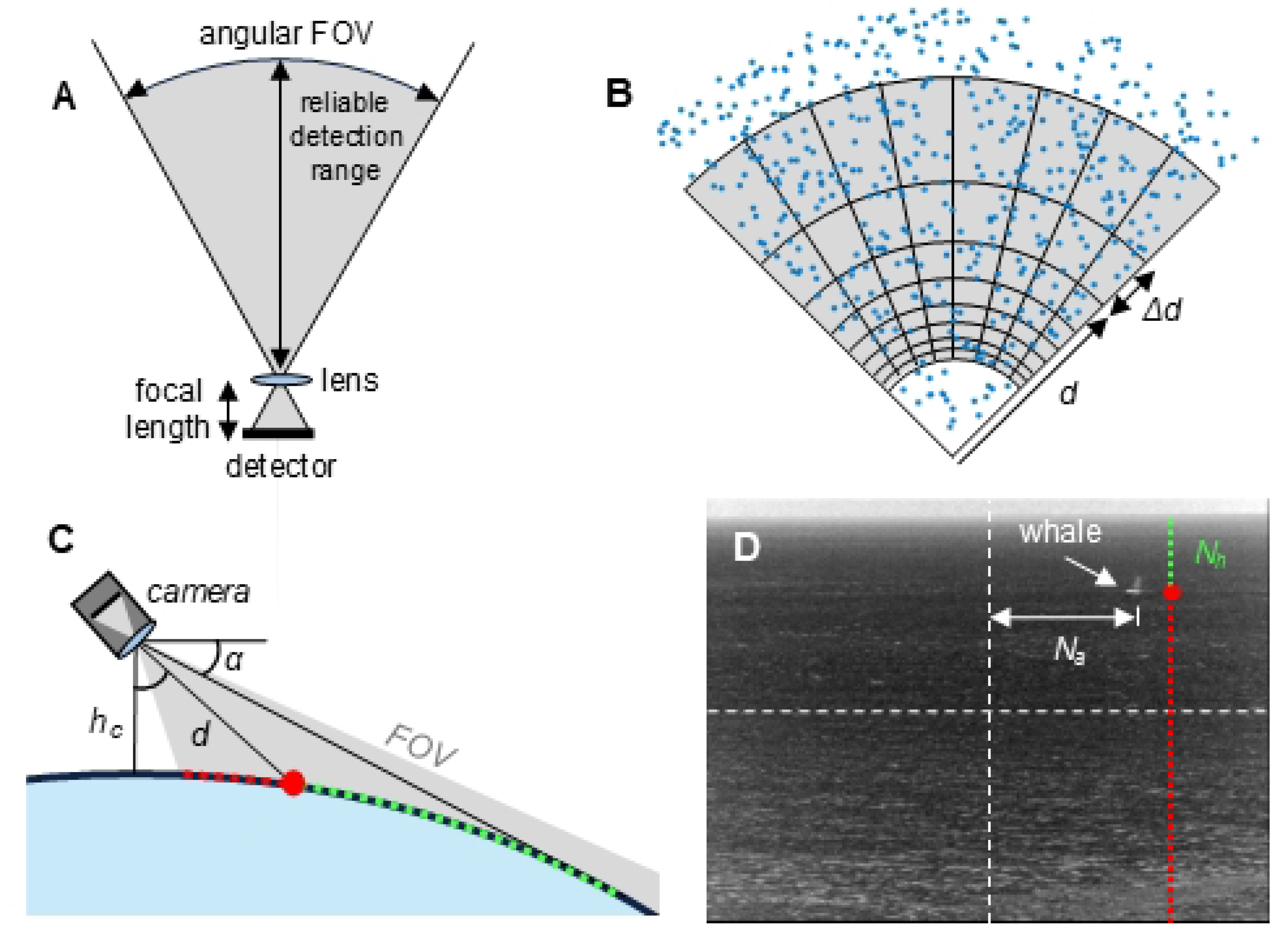
Comparison of a static camera with rotating line cameras.

## Discussion

The IR model described in this paper captures the key trade-offs and challenges in detecting cetaceans using IR systems. In normal operating conditions, the most important parameters that affect the RDR are the camera height and angular resolution (i.e., the spatial sampling of the cetacean image). For a single camera, the angular resolution is improved by using longer focal length lenses, which has the negative impact of reducing the FOV. Therefore, successful implementation of IR systems for preventing harm to cetaceans depends heavily on determining the FOV and RDR required for detecting cetaceans at distances far enough to allow for mitigation actions. To determine these requirements for reducing vessel strikes of cetaceans such as large whales, numerous factors need to be modeled in addition to the IR system’s detection capabilities. These include the IR system pipeline (e.g., system latency, data communication type, human analyst review time), vessel parameters (e.g., speed, size), and whale behavior (e.g., dive characteristics, blow interval, speed) (11).

To overcome the trade-off of FOV and angular resolution, multiple static cameras or rotating line cameras can be used. Line scanning cameras have increased FOR, but typically have lower frame rates, which reduces the RDR (Fig 9). Additional effects of rotating line scanners such as distorted frames caused by exceeded gimbal capacity during a single frame are not included in the model and may further reduce the RDR. Multiple cameras can maintain high frame rates but have increased data throughput which increases processing time and complicates detection algorithms. One alternative may be fast scanning of a small FOV camera to a region of interest identified with a large FOV camera, as implemented by Tilmon et al. with a micro-electro-mechanical system (MEMS) mirror (25).

Although the focus of the model is on RDR, the minimum detection range must also be considered when decreasing the FOV. As the FOV becomes smaller, the angle of the camera relative to sea level should be optimized to detect cetaceans over the range of interest.

Using the onshore detection data from Guazzo et al., we were able to validate the performance of the developed model for this IR detection scenario. Because fewer cetaceans are closer to shore and were localized to a migration route, we were concerned that a uniform whale density over the detection range of the IR system could be a poor assumption. Indeed, the number of whales predicted with the model at short detection distances is higher than the number of whales detected in the study (Fig 5B and 5D). Fortunately, Guazzo et al. state that the visual sightings decreased at a lower rate than the IR detections as the distance increased, indicating the decrease in whale detections by the IR system at larger distances was due to IR system limitations and not a decrease in whale density.

To be clear, the RDR calculated in the model does not depend on the assumption of uniform whale distribution, only the histogram of cetaceans as a function of distance does. This assumption may not be valid in many cases because cetaceans aggregate and the distribution over the sea is nonuniform. The empirical method for calculating RDR currently depends on making this assumption: detections of cetaceans are plotted in a histogram, and the peak of the histogram is assumed to be the RDR, which is only valid if the distribution is uniform. Therefore, model validation using data collected by IR systems can be improved by factoring in the expected distribution of cetaceans over the region of detections.

The large number of detections from the Guazzo study (1882 whales), clear conditions, and the stability of the onshore mount provided a robust dataset for validation, but we were unable to obtain any additional dataset that could be used for testing. This is a significant limitation for the work because testing additional parameters is critical for ensuring the accuracy of the model. One potential way to implement this model would be to include a small dataset for an IR system of interest. This dataset could validate the model and provide any calibration inputs that may be required. The model could then simulate a wide range of operating conditions to determine the IR system’s expected performance.

Although many of the key parameters are captured in the model, it should be further developed to include effects of Beaufort scale, gimbal capacity, visibility as a function of the wavelength range for the IR camera (which would help capture differences in the effects of humidity), and additional components missing in how the detection algorithm works (9). In cases for which the gimbal capacity is exceeded for a vessel at sea, the image of the blow may become distorted and difficult to detect (1). Further, higher winds produce white caps that could increase false positives in the detection algorithm, as well as disperse the blow in a shorter amount of time (12,23). All these effects would make it more challenging to detect cetaceans over far ranges, so the model in its current state may overestimate the RDR.

## Conclusion

IR imaging systems are an emerging tool for mitigating the risk of vessel-whale collision for both manned and autonomous vessels and are also used to help mitigate risks to whales from offshore construction and survey activities. Developing models for these various settings with IR systems that have a massive parameter space is a challenging problem to solve. The model developed here is a helpful step towards evaluating the efficacy of IR systems used in these applications and may help inform the critical parameters required to achieve mitigative actions.

## Acknowledgments

The authors would like to acknowledge Trevor Joyce, Eric Patterson, and Caroline Good for providing IR camera data and feedback for the manuscript.

This software and technical data were produced for the U. S. Government under Contract Number 1331L523D130S0003, and is subject to Federal Acquisition Regulation Clause 52.227-14, Rights in Data—General, Alt. II, III and IV (DEC 2007) [Reference 27.409(a)].

No other use other than that granted to the U. S. Government, or to those acting on behalf of the U. S. Government under that Clause is authorized without the express written permission of The MITRE Corporation.

For further information, please contact The MITRE Corporation, Contracts Management Office, 7515 Colshire Drive, McLean, VA 22102-7539, (703) 983-6000. **© 2026 The MITRE Corporation**. Approved for Public Release; Distribution Unlimited. Public Release Case Number 25-3252.

